# Cyclic tri-adenylate signalling by a Panoptes anti-phage guard system with a CARF-TM effector

**DOI:** 10.64898/2026.03.13.711614

**Authors:** Sabine Grüschow, Peter Wotherspoon, Emma Hilton-Balfe, Shirley Graham, Malcolm F White

## Abstract

Cyclic nucleotide second messengers are used in all domains of life to amplify viral infection signals and activate cellular defences. In prokaryotes, CBASS (cyclic nucleotide based antiphage signalling system) and type III CRISPR-Cas systems generate a range of cyclic nucleotides which bind and allosterically activate effector proteins to mount an anti-viral response. Viruses have evolved counter-measures to antagonise these signalling pathways in the form of cyclic nucleotide sponges and phosphodiesterases that sequester or degrade these molecules to subvert immunity. Recently, the Panoptes system was shown to function as a guard against these viral tactics. The type I Panoptes polymerase, mCpol, generates cyclic dinucleotides as decoy molecules that, when sequestered by phage proteins, results in the activation of the membrane-permeabilising effector 2TMβ to halt the phage infection cycle. Here, we investigate the type II Panoptes system, demonstrating that it generates cyclic tri-adenylate (cA_3_) to maintain a CRISPR-associated Rossmann fold-transmembrane (CARF-TM) effector in an inactive, dimeric state. When cA_3_ is sequestered or degraded, the CARF-TM protein oligomerises, resulting in increased outer membrane permeability and growth arrest. These findings expand our understanding of the guard systems that constitute a fascinating component of the bacterial immune system.

## Introduction

In recent years, a large number of antiviral defence systems have been discovered in prokaryotes, leading to the concept of the bacterial immune system [1, 2]. One major approach, used in both prokaryotes and eukaryotes, is the generation of nucleotide second messengers by specialised nucleotide cyclase enzymes activated by viral infection [3, 4]. Notable examples include the cGAS/STING pathway in metazoa which signals via cGAMP [5] and its bacterial equivalent CBASS (cyclic nucleotide based antiphage signalling system) [6], Pycsar (Pyrimidine cyclase system for antiphage resistance) which generates cyclic pyrimidines [7] and the type III CRISPR-Cas system signalling via production of cyclic oligoadenylate (cOA) species [8, 9]. This approach has the advantage of achieving amplification of the primary signal of infection – for example the type III CRISPR system detects 1 viral mRNA molecule and generates over 1000 cyclic nucleotides in response [10]. Viruses, in turn, target the cyclic nucleotides to neutralise cellular defences using specialised phosphodiesterases that degrade cGAMP in eukaryotes [11] and bacteria [12], ring nucleases that target cOA species to nullify CRISPR-based immunity [13] and sponge proteins that bind cyclic nucleotide second messengers [14, 15]. Sponge proteins are particularly adept at sequestering a range of cyclic nucleotides used by immune defence systems and are widely encoded by phage [16, 17].

Type III CRISPR systems use their catalytic Cas10 subunit, which has two Palm polymerase domains, to generate cOA species from ATP [8, 9]. The ring size of cOA species produced ranges from 3 to 6 AMP subunits, activating a diverse range of effector proteins to provide an immune response [18, 19]. The recent discovery of variants that make the signalling molecule SAM-AMP by conjugating S-adenosyl methionine and ATP [20] underscores the flexibility of the polymerase to accept a range of building blocks for signal generation. In 2015, Burroughs et al. described a minimal CRISPR-related polymerase (mCpol), distantly related to Cas10, which was sometimes associated with CARF domain proteins characteristic of type III CRISPR effectors [21]. In some cases, the mCpol cyclase is fused to a Csx1-family ribonuclease effector protein, potentially representing a single-protein antiviral defence system that functions via cyclic nucleotide signalling [22]. In other cases, 2 gene operons include mCpol next to a gene encoding a predicted membrane-bound effector [22]. mCpol has been suggested as the ancestral protein that gave rise to Cas10 and the evolution of the class 1 CRISPR systems [22].

Recently, two research groups independently demonstrated that one type of mCpol-containing system, which they named “Panoptes” (henceforth Panoptes type I, Fig. 1A), generates cyclic dinucleotides constitutively, maintaining an associated 2TMβ (2 trans-membrane helix, β-strand rich) effector in an inactive state [23, 24]. This contrasts with CBASS, and other defence systems relying on cyclic oligonucleotide signalling, which requires an active phage infection to trigger cyclic nucleotide synthesis. When a phage expressing an anti-defence sponge protein such as Acb2 infects cells encoding Panoptes type I, the cyclic dinucleotide is depleted, resulting in activation of the 2TMβ-effector, leading to growth arrest and thwarting the phage infection [23, 24] (Fig. 1B). Panoptes is thus an example of a guard system that may often work in conjunction with CBASS defence to prevent phage from utilising a cyclic nucleotide depletion strategy.

**Fig. 1.**
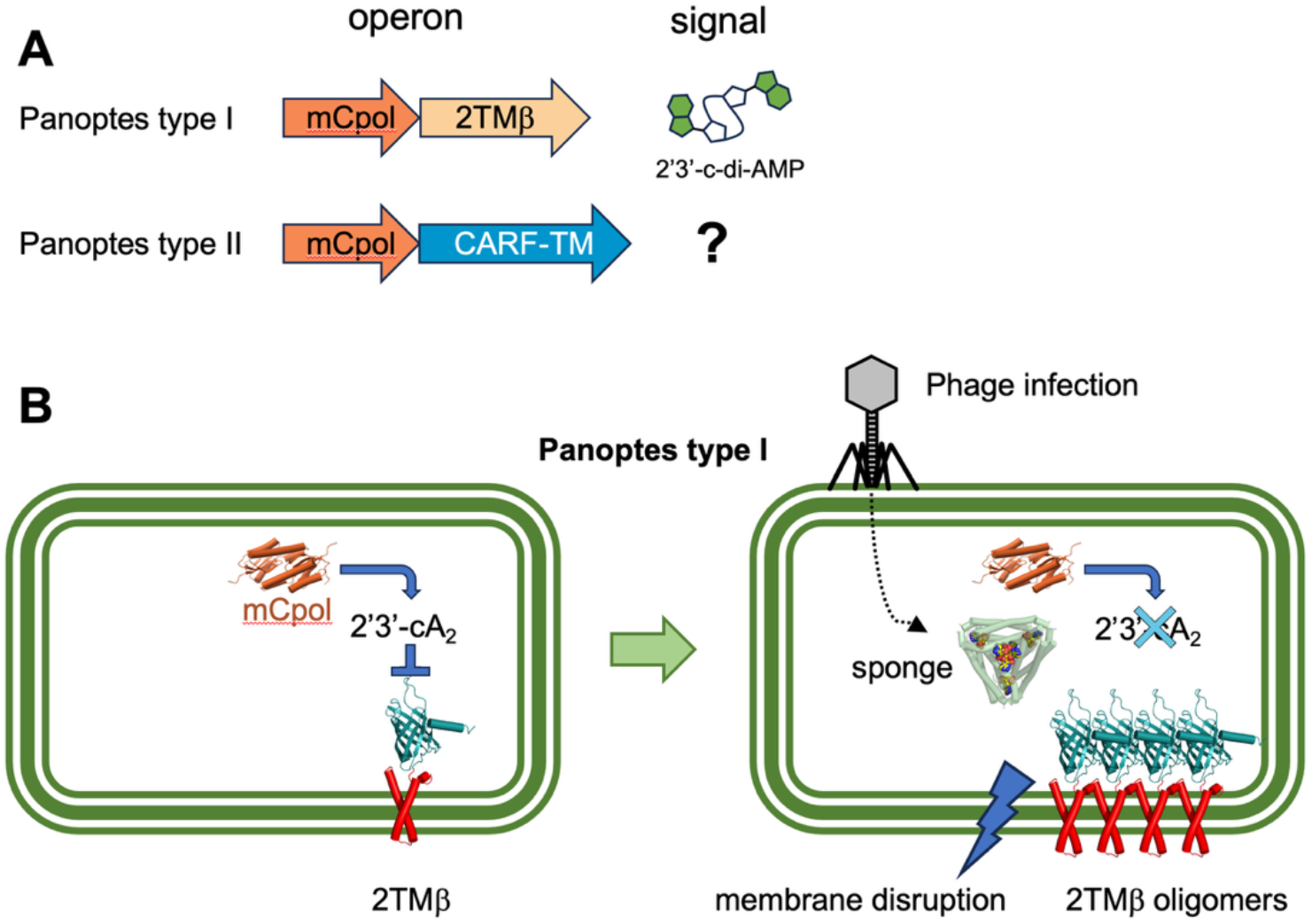
Organisation and mechanism of the Panoptes System. **(A)** Operon organisation and major cyclic nucleotide signal for representative type I and II Panoptes systems. **(B)** In type I Panoptes systems, mCpol is constitutively active, generating signalling molecules that inactivate the cognate effector, 2TMβ. When an infecting phage expresses a sponge protein, the cyclic oligonucleotide pool is removed. The Panoptes effector oligomerises, leading to membrane disruption and growth arrest or cell death.

Here, we investigate a different type of Panoptes system (Panoptes type II) that employs a CARF-family effector with a transmembrane domain (CARF-TM). The mCpol cyclase generates a cyclic trinucleotide (cA_3_) product which is bound by the associated CARF-TM effector. We demonstrate that that the effector is dimeric in the presence of cA_3_. The absence or removal of cA_3_ from CARF-TM results in structural changes that ultimately lead to outer membrane (OM) permeabilization and render cells incapable of forming colonies. These data demonstrate another facet of Panoptes-mediated immunity and the potential to guard type III CRISPR-Cas as well as CBASS defence systems.

## Results

### Type II Panoptes signals via cyclic tri-adenylate (cA_**3**_)

While the characterised type I Panoptes systems synthesise cyclic dinucleotides and utilise a 2TMβ - family effector, type II systems with a CARF-TM effector remain unstudied. We designed synthetic genes encoding mCpol and CARF-TM from the cyanobacterium *Cylindrospernum stagnale* PCC7417 (Cst) for expression in *E. coli* and purified the recombinant wild-type and variant proteins. The Cst mCpol protein was tested for nucleotide cyclase activity and shown to synthesise cA_3_ in the presence of ATP and Mg^2+^ (Fig. 2). The presence of other NTPs did not alter the reaction product. The turnover number was approximately 1 h^-1^ *in vitro* (Fig. S1), making mCpol significantly slower than Cas10 from Type III CRISPR systems [25]. A second mCpol associated with a CARF-TM effector, from *Sphingomonas* (Sph), also synthesised cA_3_, also with a slow turnover number of 1 h^-1^, suggesting that this is likely to be a common feature of the CARF-TM associated systems. Residues D11 and D63 of Cst mCpol correspond to the canonical metal binding residues of Cas10 and Palm family polymerases more generally. We mutated D63 and the neighbouring D64 to generate Cst mCpol[D63N/D64N]. This variant was completely inactive, as expected (Fig. 2).

**Fig. 2.**
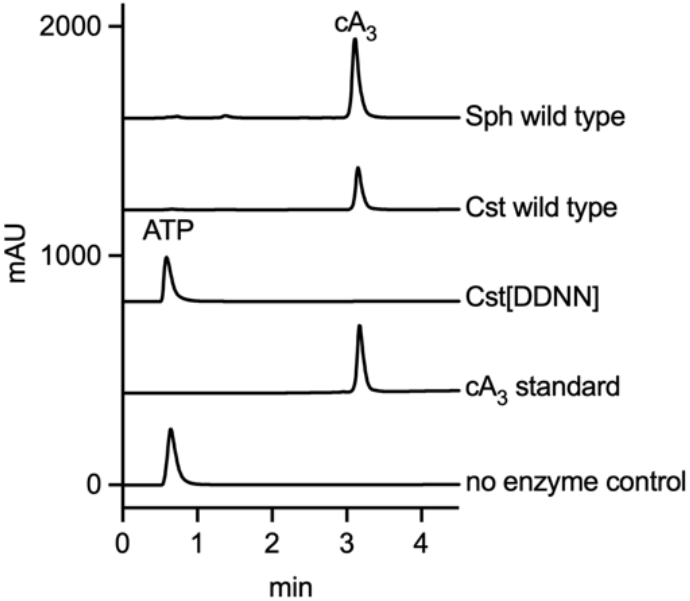
Type II mCpols synthesis cA_3_. HPLC analysis of mCpol reaction products showing production of cA_3_ in the presence of ATP. The Cst mCpol[D63N, D64N] variant (Cst[DDNN]) was catalytically inactive.

### Type II Panoptes is active *in vivo* and responds to cA_**3**_ levels in cells

To test the activity of Panoptes type II *in vivo*, we performed transformation assays, where the effector was introduced to a strain containing the mCpol cyclase. A serial dilution of the transformation mixture was then plated onto selective medium and both cyclase and effector expression were induced. The plates were incubated overnight and inspected for colony growth. The system was deemed active if fewer colonies were observed compared to controls. We used Cst mCpol protein as the cyclase with either its cognate CARF-TM effector, or the cA_3_-activated NucC protein, which is associated with CBASS and type III CRISPR defence [26, 27]. In this experiment, mCpol was expressed under control of the T7 promoter (IPTG inducible), while NucC or CARF-TM were expressed under control of the arabinose-inducible pBAD promoter. As expected, expression of NucC alone resulted in efficient transformation, as this enzyme is inactive in the absence of cA_3_ (Fig. 3A). In contrast, expression of the CARF-TM effector alone resulted in toxicity and therefore no viable colonies in the absence of mCpol. When the NucC plasmid was transformed into cells expressing Cst mCpol, no transformants were observed, consistent with activation of the nuclease by cA_3_. In contrast, the toxicity of the CARF-TM effector was moderated with co-expressed mCpol. Truncation of the CARF-TM effector to remove the predicted transmembrane domain completely abolished activity, demonstrating that the transmembrane (TM) domain is essential. Expression of the inactive D63N, D64N variant of Cst mCpol did not alleviate the toxicity of CARF-TM, confirming the requirement for cA_3_ synthesis.

**Fig. 3.**
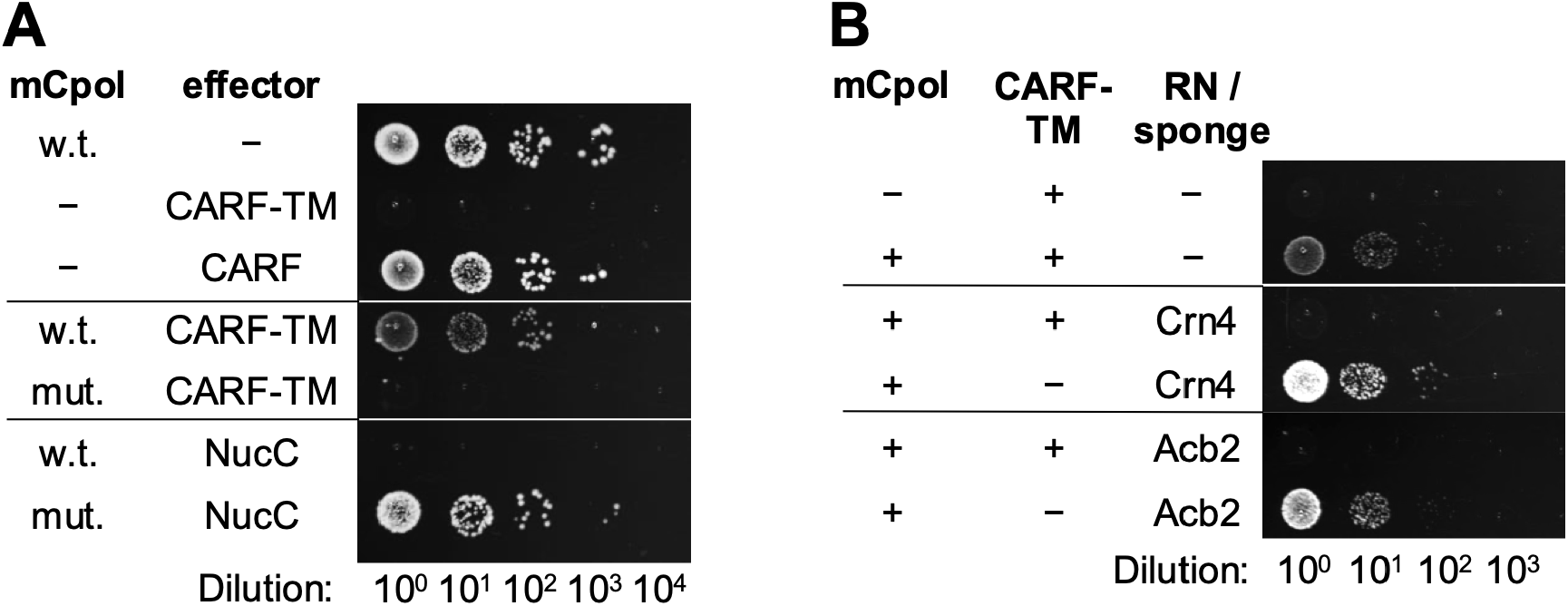
Panoptes type II signals via cA_3_ to deactivate the CARF-TM effector. **(A)** The cA_3_-activated nuclease NucC prevented cell growth only in the presence of mCpol, whereas CARF-TM did so in its absence. The transmembrane (TM) domain of CARF-TM was required for activity. w.t.: wild type; mut.: catalytically inactive mCpol[D63N, D64N]. **(B)** The presence of a cA_3_-degrading ring nuclease (Crn4) or a phage sponge protein (Acb2) reactivated CARF-TM activity.

Overall, these data establish Panoptes type II as a defence system that signals via cA_3_, using a signalling logic similar to Panoptes type I. To explore this further, we challenged the defence system with two different proteins that are known to bind or degrade cA_3_: the phage sponge protein Acb2 [14] and the cA_3_-degrading ring nuclease Crn4 [28]. Expression of either of these proteins from a plasmid resulted in the activation of Panoptes type II, reflected in cell toxicity when the CARF-TM effector was induced by arabinose (Fig. 3B). This mimics the situation where a phage utilising a sponge or ring nuclease to subvert CBASS or type III CRISPR defence would activate Panoptes type II defence.

### The CARF-TM effector binds cA_3_

Modelling with Alphafold3 [29] strongly supported a dimeric structure for the CARF-TM effector (Fig. 4A), consistent with the dimeric organisation of many other CRISPR-associated CARF effectors [22]. To express and purify recombinant CARF-TM, we had to co-express the protein with mCpol, generating cA_3_ to keep the effector in an inactive state. Under these conditions, CARF-TM with a non-cleavable N-terminal polyhistidine tag could be purified to homogeneity via immobilised metal affinity and size exclusion chromatography in the presence of the detergent DDM (Fig. S2). We also purified a truncated form of the effector (CARF), where the TM domain was replaced with a non-cleavable C-terminal hexahistidine tag. The truncated protein also required the presence of the cyclase for efficient purification. We observed two bands in the final purified sample for full length CARF-TM and for the truncated CARF protein (Fig. S2); both were confirmed by mass spectrometry to be the effector. An elevated absorbance at 260 nm for both the full length and truncated proteins was suggestive of the presence of nucleic acid. We extracted nucleotides from both proteins and analysed them by HPLC, confirming the presence of cA_3_ in roughly equimolar amounts to the protein dimer (Fig. 4C).

**Fig. 4.**
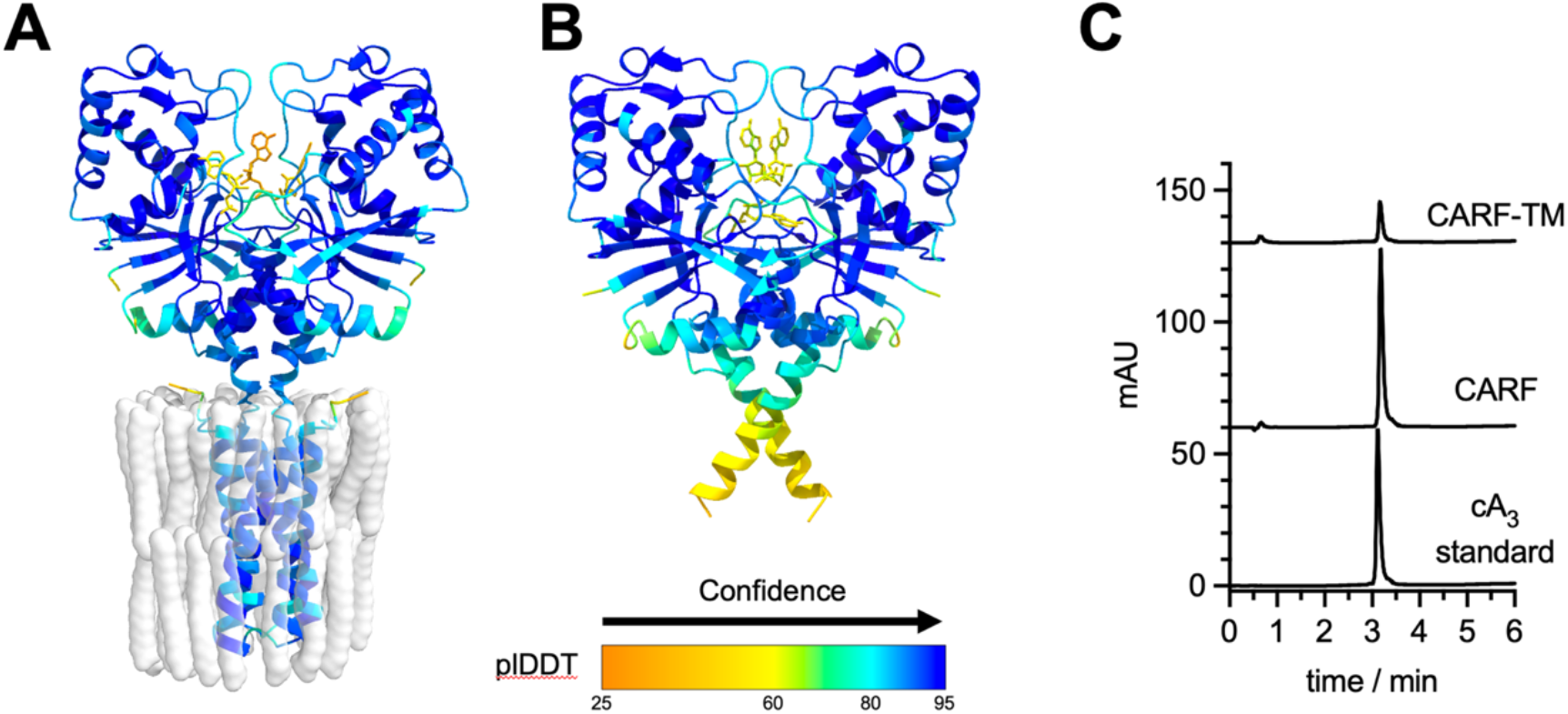
CARF-TM structure and cA_3_ binding. Alphafold3 models of CARF-TM **(A)** and the truncated CARF **(B**), coloured by plDDT value. Both were modelled as a dimer with three AMP ligands (shown as yellow sticks) to indicate the likely cA_3_ binding site. CARF-TM modelling included 50 oleic acids (shown as translucent spheres) to simulate the lipid bilayer. **(C)** HPLC analysis of the supernatant of heat denatured CARF-TM and CARF confirmed the presence of cA_3_ in the purified proteins.

These observations demonstrated that cA_3_ remains tightly bound to both the full length and truncated effector during purification, suggesting a low dissociation constant for cA_3_ binding. To explore ligand specificity, we investigated binding of ^32^P-radiolabelled cA_3_ by electrophoretic mobility shift assay (EMSA). At high concentrations of CARF-TM or CARF domain (approximately 1 µM dimer), a single retarded species consistent with cA_3_ binding became visible (Fig. 5A). This high apparent dissociation constant likely reflects the fact that both proteins were already bound to unlabelled cA_3_, requiring displacement by competition with the radioactive species. The retarded radioactive cA_3_ moiety was abolished when a 100-fold molar excess of cold cA_3_ was added, but binding was unaffected by a range of other cyclic nucleotide species (Fig. 5B). Overall, these data demonstrate that both the CARF-TM effector and its truncated, soluble CARF domain bind cA_3_ specifically and tightly, but not irreversibly.

**Fig. 5.**
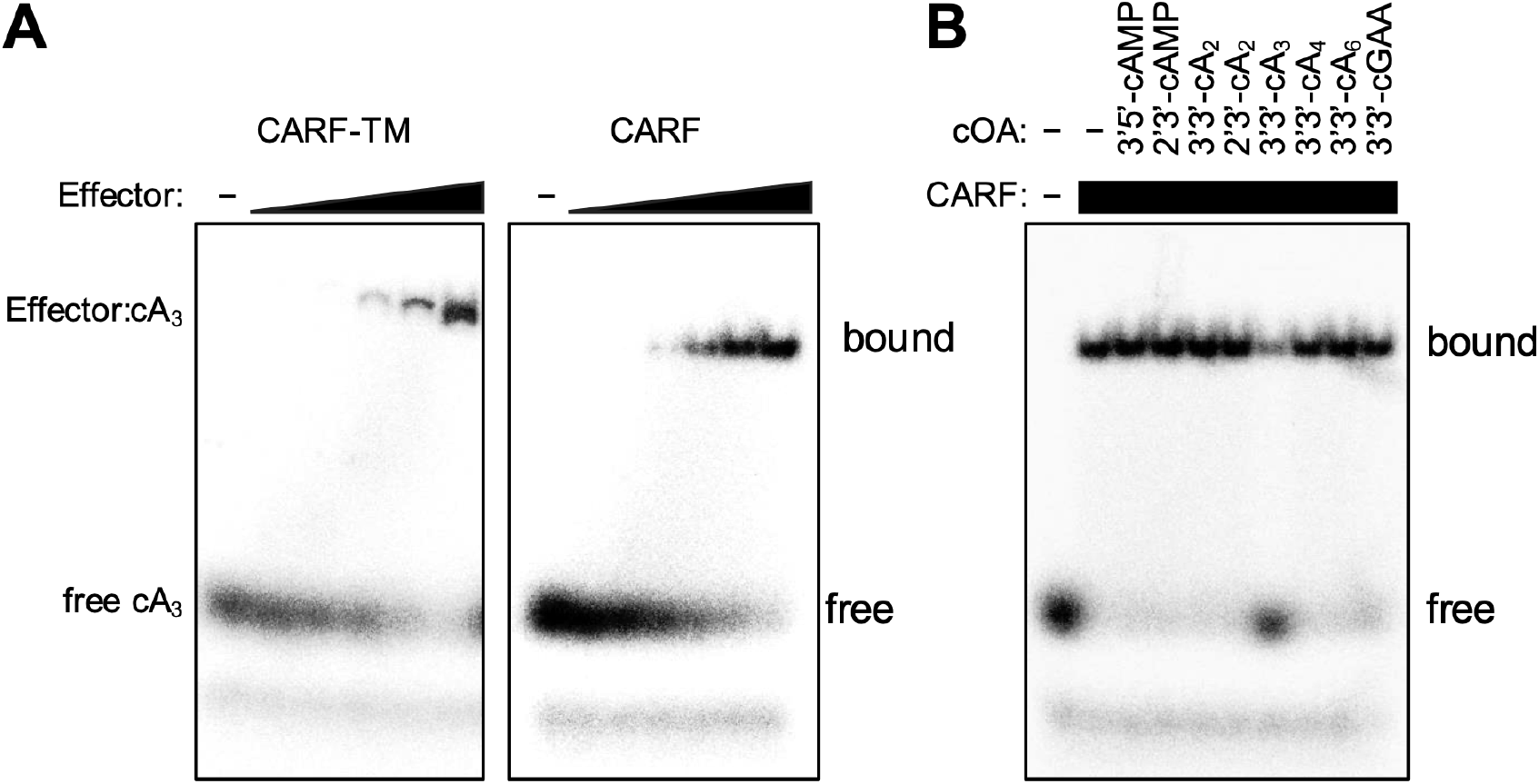
The full length and truncated CARF proteins bind cA_3_. **(A)** Radiolabelled cA_3_ (100 nM) was incubated with increasing CARF-TM (125 nM – 2 µM dimer) or CARF (200 nM – 7 µM dimer) and analysed by EMSA. **(B)** The CARF: [^32^P]-cA_3_ complex was challenged with a 100-fold excess of a range of cyclic nucleotides, as indicated. Only cA_3_ was able to displace the radiolabelled bound ligand.

### The soluble CARF variant multimerises when cA_**3**_ is not bound

We next investigated whether the CARF effector underwent any conformational changes upon loss of cA_3_, mimicking the effect of a cA_3_-scavenging phage protein. Although we could purify small quantities of the full length CARF-TM effector, plasmid challenge experiments indicated that the presence of N- or C-terminal tags abolished function *in vivo* (Fig. S3). We therefore focused on the truncated CARF version of the effector for further *in vitro* studies. To deplete cA_3_ from the CARF protein, we incubated the purified protein with increasing concentrations of the ring nuclease Crn4. HPLC analysis of the nucleotide species present following Crn4 incubation confirmed that a significant proportion of the cA_3_ present was converted to smaller products, confirming that the ring nuclease can capture and partially degrade CARF-bound cA_3_, resulting in the generation of *apo*-protein (Fig. S4).

To examine the effect of cA_3_ removal on the effector, we probed structural changes using the fluorogenic dyes Sypro Orange and BODIPY FL L-cystine (BFC). Both dyes are typically used with thermal shift assays; however, we conducted isothermal assays following changes in fluorescence after addition of Crn4 to the cA_3_:CARF complex. The fluorescence intensity of Sypro Orange increases upon binding to hydrophobic regions, which are normally buried in a folded protein but become exposed when the protein denatures and in turn become hidden when an unfolded protein aggregates (Fig. 6A). The fluorescence of BFC in its disulfide dimer form is quenched. Cleavage by free thiols, such as exposed cysteine residues, releases the monomeric, fluorescent dye (Fig. 6B). Unfolding and / or increased protein chain mobility can expose buried cysteine residues leading to an increase in fluorescence.

**Fig. 6.**
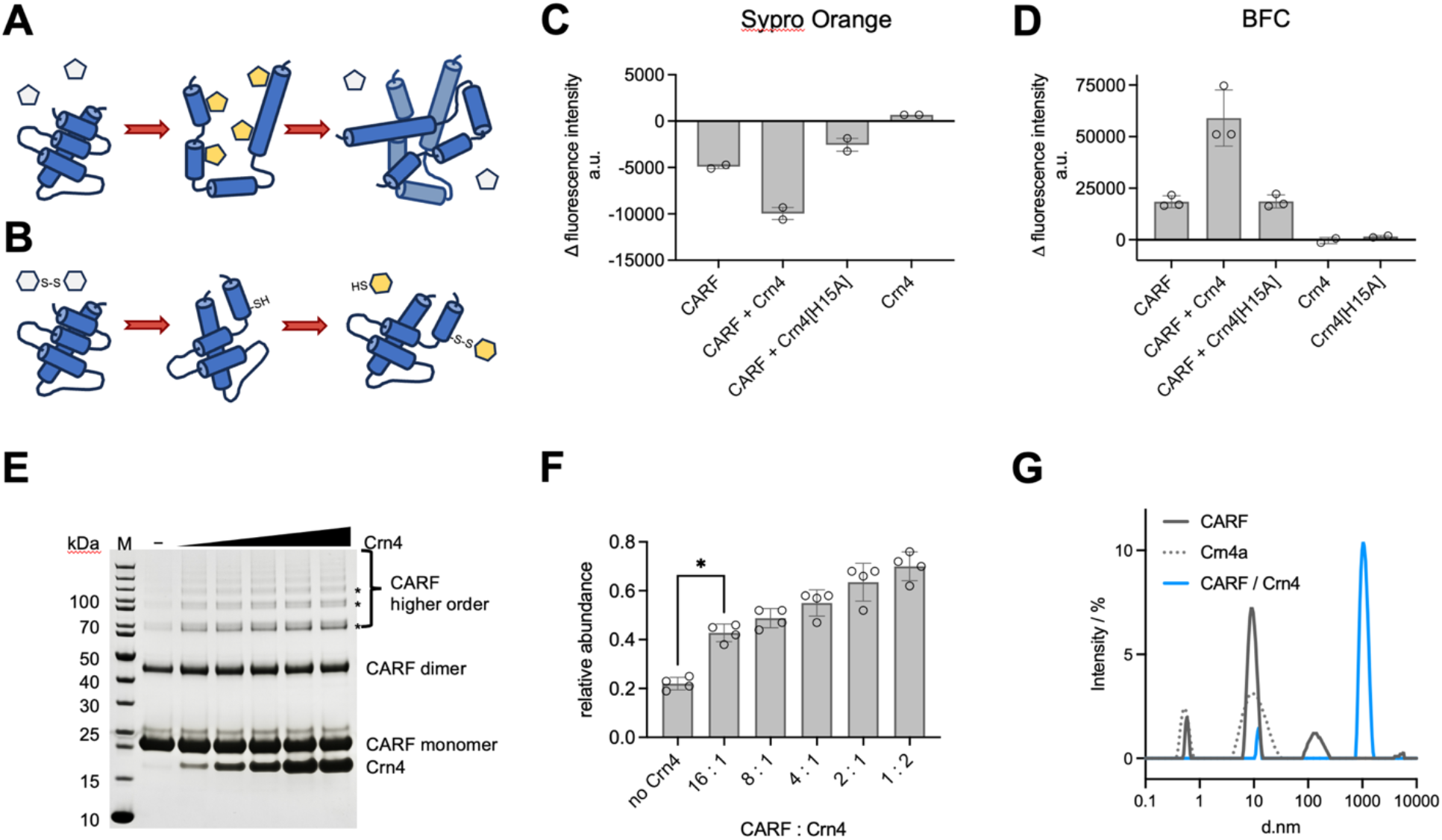
Removal of cA_3_ results in oligomerisation of CARF-TM. **(A)** Cartoon of Sypro Orange binding to exposed apolar patches upon protein unfolding followed by occlusion of hydrophobic binding sites through protein aggregation. (**B**) The BFC dimer becomes fluorescent upon cleavage of its disulfide bridge by cysteine residues. (**C, D**) Difference in fluorescence intensity after 50 min incubation of CARF with Crn4 wild type or catalytically inactive H15A variant. The greatest change in fluorescence was observed for CARF with wild type Crn4. Removal of cA_3_ appears to remove available hydrophobic residues (**C**, Sypro Orange, mean and range of duplicates shown) but expose Cys residues (**D**, BFC, mean and standard deviation of triplicates shown). (**E, F**) Chemical cross-linking of CARF in the presence of increasing amounts of Crn4. CARF and Crn4 were incubated for 30 min before separating on PAGE (**E**). CARF oligomers labelled with an asterisk in E were quantified by densitometry relative to the CARF dimer band. **G:** DLS analysis of CARF after incubation with Crn4.

We observed a decrease in Sypro Orange fluorescence intensity upon Crn4 treatment, consistent with burial of hydrophobic patches upon ligand removal due to protein aggregation (Fig. 6C). For the BFC dye, isothermal incubation of the CARF protein with Crn4 resulted in a significantly larger increase in fluorescence than in the absence of Crn4 or with the catalytically inactive Crn4 [H15A] variant [28] (Fig. 6D).

The effect of cA_3_ removal on the quaternary structure of the CARF protein was also tested using chemical cross-linking (Fig. 6E). In the absence of Crn4, the CARF protein was observed primarily as monomeric and dimeric after crosslinking, with very minor amounts of larger species. With increasing amounts of Crn4, progressively higher amounts of large (>dimeric) cross-linked species were observed (Fig. 6F). This assay benefitted from the unusual amino acid composition of Crn4, which lacks lysine residues and thus cannot cross-link. We also investigated the removal of cA_3_ from the CARF domain by dynamic light scattering (DLS). Notably, a precipitate formed when Crn4 was incubated with the CARF domain. As the samples were centrifuged, the precipitate was removed prior to DLS analysis. An approximately 100-fold increase in size of the remaining soluble CARF protein was detected upon cA_3_-depletion (Fig. 6G).

Together, these observations suggested that cA_3_ removal results in structural perturbations of the CARF dimer, leading to unfolding or aggregation *in vitro*. Given the difficulty in studying the full length CARF-TM protein *in vitro*, we turned to *in vivo* analyses.

### The CARF-TM effector causes growth arrest in the absence of cA_3_

We set out to monitor changes in cell viability upon expression of CARF-TM using microscopy. The full length, untagged CARF-TM was expressed from the T7 promoter in pEHisV5TEV [25], allowing induction of expression using IPTG. We omitted expression of the mCpol from these strains, so that all effectors would be in the active *apo* form when expressed. *E. coli* harbouring CARF-TM, truncated CARF or the empty vector were grown to mid-log phase before induction with IPTG. Cells collected 1 - 2 h after induction were stained with Syto9 and propidium iodide (PI) [30]. Both are nucleic acid staining fluorescent dyes; Syto9 is a membrane-permeable dye that will stain live and dead cells, whereas PI can only enter cells with damaged membranes. PI has a higher affinity for nucleic acids than Syto9 and tends to displace Syto9 from the nucleic acid given a high enough concentration. We determined the percentage of PI-dyed cells using fluorescent confocal microscopy (Fig. 7A). We observed only minor differences between CARF-TM expressing (7.2% PI) or non-expressing cells (3.5% PI for empty vector, 2.3% for CARF). This suggested that CARF-TM is not a fast-acting bactericidal agent when expressed in *E. coli*.

**Fig. 7.**
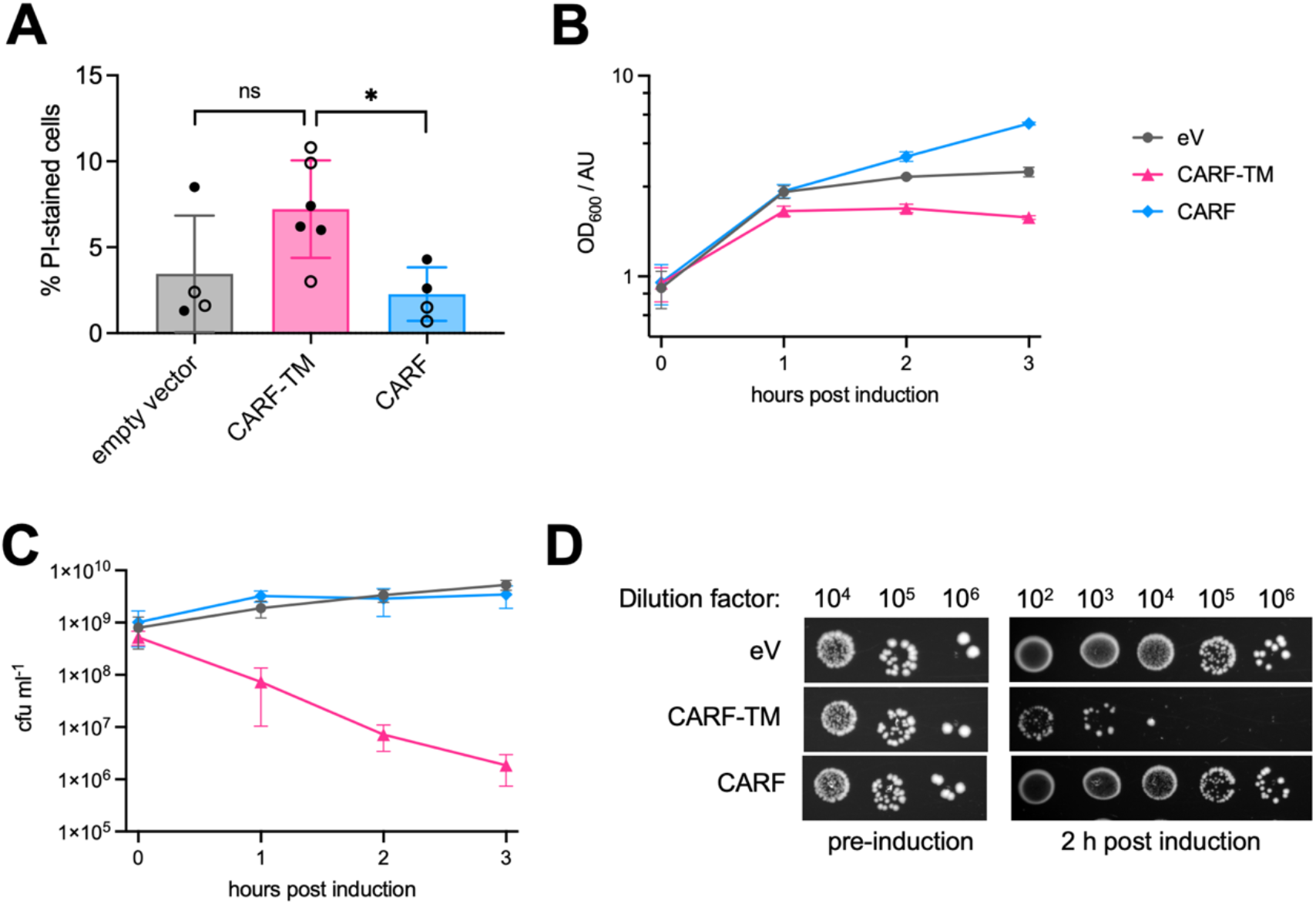
The CARF-TM effector causes growth arrest in the absence of cA_3_. (**A**) Live/dead staining of cells expressing CARF-TM, CARF or empty vector. Samples from 1 h (open circles) or 2 h (closed circles) after induction of the effector protein were analysed by confocal fluorescence microscopy. There was no significant difference in the percentage of PI-stained cells between strains expressing CARF-TM or those lacking the effector (p value 0.11, two-tailed Mann-Whitney test); however, cells expressing the truncated effector CARF had a lower percentage of PI-staining than those expressing full length CARF-TM (p value 0.019). Data show the mean and standard deviation from three independent experiments. More than 1000 cells were counted in total for each strain. (**B**) Optical density of cultures after induction of protein expression. (**C, D**) Cells expressing CARF-TM lost their ability to proliferate as determined by microdilution assay. Data show the mean and standard deviation of four biological replicates for panels B, C. One representative example of the microdilution assay is shown in D. eV: empty vector.

To follow on from the live/dead staining, we determined the cell density and culturability of the three strains for the first three hours after induction of CARF-TM expression. The optical density initially increased for all three strains, levelling off for the CARF-TM-expressing strain after 2 h, consistent with growth arrest upon effector expression (Fig. 7B). Microdilution and colony counting assays demonstrated that whilst the strains with empty vector or expressing the CARF domain yielded a similar number of colony-forming units (cfu), those for the CARF-TM-expressing strain dropped by 2 – 3 orders of magnitude (Fig. 7C, D). Combined with the data from PI staining, this hinted at dormancy akin to the viable-but-not-culturable (VBNC) state [31] when the CARF-TM protein was expressed in the absence of cA_3_.

### Expression of apo CARF-TM changes cell permeability in *E. coli*

We next assessed the effect of CARF-TM on metabolic rate by monitoring the cellular reduction of resazurin to resorufin [32] by fluorescence spectrometry (Fig. 8). Before induction, the fluorescent signal was indistinguishable between the three strains (CARF-TM, CARF and eV). By 1 h after induction, CARF-TM containing cells showed a faster rate of resorufin production (suggestive of increased metabolic rate) compared to control cells (Fig. 8A). As a control, we added polymyxin B, which permeabilises the outer membrane (OM) of Gram-negative cells [33, 34]. On addition of polymyxin B to the resazurin reaction, the rate for the CARF-TM containing strain did not change significantly after induction of protein expression and was comparable to the control strains (Fig. 8B). This suggested that OM permeabilization, similar to that observed by polymyxin B action, might be responsible for the apparent increase in NAD(P)H levels upon CARF-TM expression. To explore this further, we investigated cell staining using the lipophilic dye Nile Red [35]. Cells expressing the apo CARF-TM effector were readily stained by Nile Red (Fig. 9). Under the same conditions, only a small fraction of cells expressing the CARF domain or containing the empty vector were stained with Nile Red (Fig. 9). These data reinforce the conclusion that expression of apo CARF-TM results in increased OM permeability. Together, these data support a model where expression of apo CARF-TM in the inner membrane of *E. coli* results in permeabilization of the OM and growth arrest, but not cell death.

**Fig. 8.**
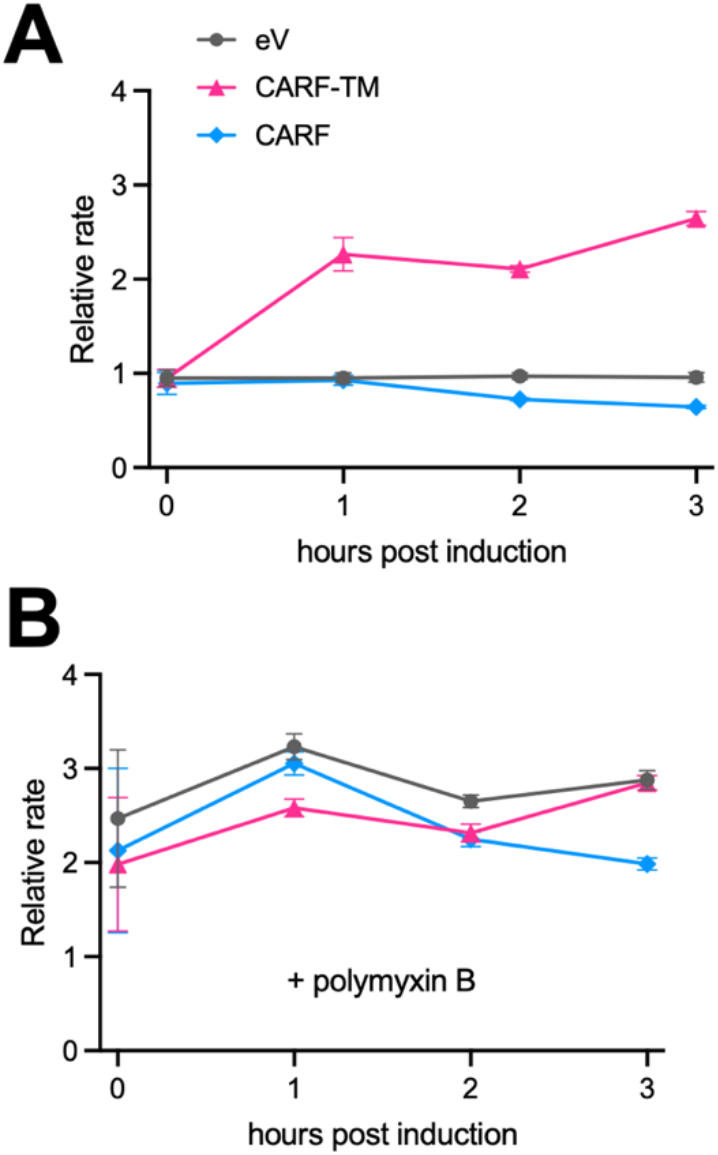
Expression of apo CARF-TM perturbs metabolic activity. Initial rates of resazurin reduction in the absence (A) and presence (B) of polymyxin B relative to empty vector (eV). Data show the mean and standard deviation of four biological replicates.

**Fig. 9.**
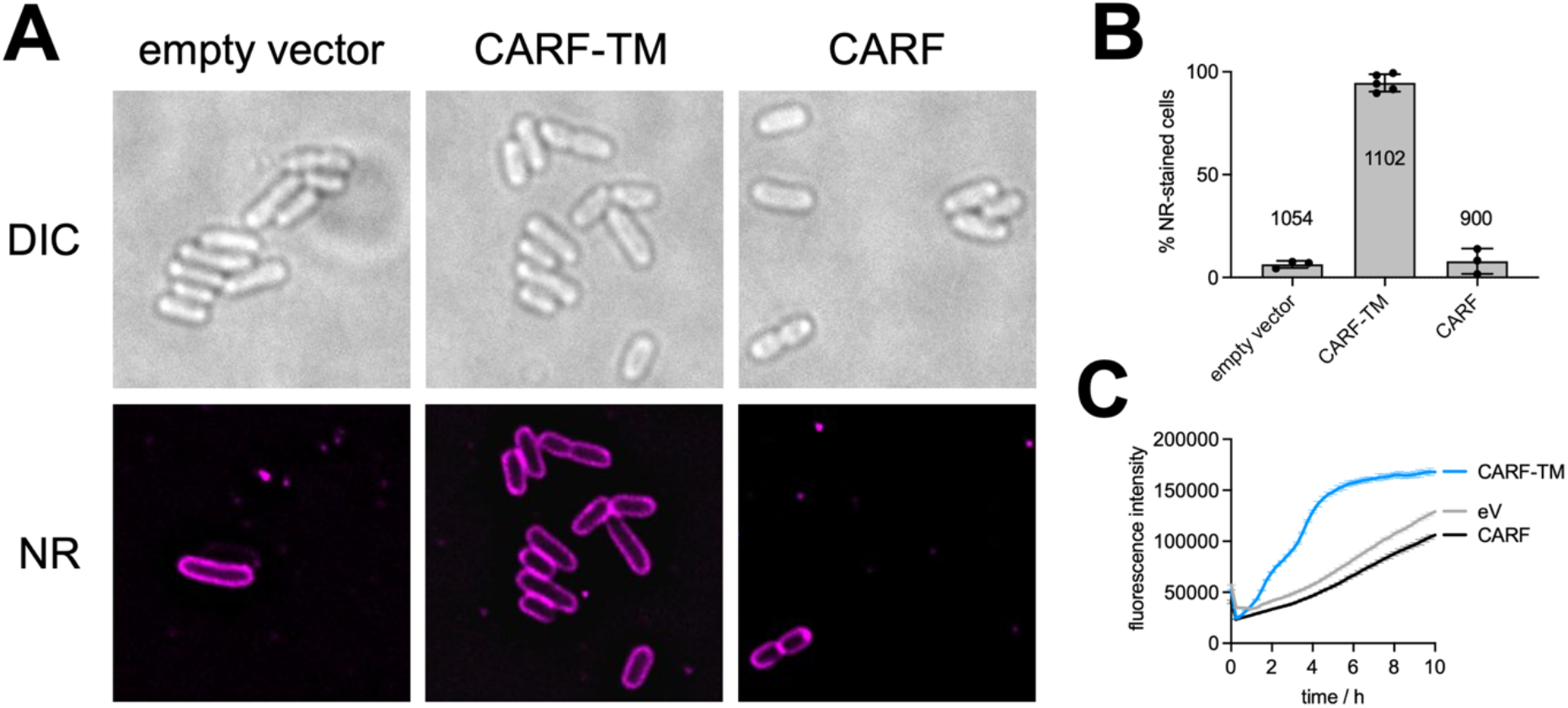
The apo form of the CARF-TM effector alters membrane permeability. **(A)** Differential interference contrast (DIC) and fluorescence microscopy of *E*. coli cells expressing the Panoptes effector CARF-TM. Cells were collected one hour after addition of IPTG to induce expression of the effector protein and stained with Nile Red (NR). Membranes of *E. coli* expressing CARF-TM were readily stained with Nile Red, whereas most cells expressing the truncated CARF domain or carrying the empty vector remained unstained. (**B)** Percentage of NR-stained cells between 1 and 2 h post induction. The total number of cells for each strain is indicated in the graph. Data are from at least two independent experiments. (**C)** NR-fluorescence is increased upon expression of CARF-TM relative to CARF or empty vector in liquid culture.

## Discussion

Here, we have shown that type II Panoptes systems with CARF-TM effectors, which constitute around 40% of total instances [23], signal using cyclic triadenylate (cA_3_). To function as a guard system, Panoptes must generate a cyclic nucleotide that is not used by the other defence systems present, but which is targeted by phage for sequestration or degradation. Type II Panoptes could co-exist and cooperate with the majority of type III CRISPR systems, which signal via cA_4_ or cA_6_ [19], and with the majority of CBASS systems, which utilise cyclic di- or tri-nucleotides [36]. A comprehensive analysis of the distribution and gene linkage of mCpol revealed that about 50% of Panoptes operons are found in genomes that also encode CBASS defence, occurring in the same gene neighbourhood in half of these instances [23]. A significant enrichment of type III CRISPR systems in Panoptes-containing genomes was also noted [23]. The *C. stagnale* genome, harbouring a type II Panoptes system, also encodes a type II CBASS system with a Cap4 effector closely related to enzymes that are activated by 2’3’3’-cA_3_ and 3’3’3’-cAAG [37, 38]. Likewise, the Sph mCpol gene is situated adjacent to a type II CBASS system with a Cap4 effector. Both of these mCpol systems could thus be using 3’3’3’-cA_3_ as a decoy for the CBASS signalling molecules. On the other hand, the Cst genome also encodes two type III CRISPR systems which appear to function via cA_4_ signalling, with Csx1 and Can2 effectors [19] – a situation amenable to the deployment of a cA_3_ decoy molecule.

The use of cA_3_ as a guard molecule in turn suggests that phage target this molecule to circumvent CBASS or CRISPR defence. This has not yet been observed directly *in vivo*, however phage-encoded Acb2 sponge proteins bind cA_3_ *in vitro* [14]. Recently, a ring nuclease, Crn4, was shown to degrade cA_3_ and to be encoded in some phage genomes [28]. We confirmed functional activation of the type II Panoptes system by both Acb2 and Crn4 *in vivo* (Fig. 3).

Activation of type I Panoptes with a 2TMβ effector results in inner membrane (IM) collapse and cell death [24]. For type II Panoptes, all our observations point to permeabilization of the OM upon expression of CARF-TM in apo form (mimicking the situation where a phage infection has disrupted cA_3_ signalling) and growth arrest, but not cell lysis. We can be confident that the CARF-TM protein is situated in the IM with the CARF domain facing the cytoplasm where cA_3_ is synthesised. The inner and outer membranes are of course functionally linked [39]. OM phospholipids, lipopolysaccharides and outer membrane proteins (OMPs) [40] are synthesised in the cytoplasm and transported through the IM to the OM [41]. Changes in any of the three OM components have the capacity to cause OM disruptions [42]. Disruption or depolarisation of the IM can thus have knock-on effects on the OM. Further investigation will be required to differentiate causality and indirect effects, ideally using a cognate host and genomic expression of the Panoptes system from native promoters.

Rapid advances in recent years have resulted in the identification of hundreds of prokaryotic antiviral defence systems [1, 2]. The “average” bacterium encodes over 7 different systems, but it has not been clear whether they often function synergistically to provide antiviral immunity. Guard systems such as Panoptes, and the Hailong system that operates via a decoy ssDNA molecule [43], are an intriguing facet of bacterial anti-phage immunity, providing the opportunity for synergistic interactions between different defence components [23]. Undoubtedly, further examples will be uncovered in the coming years.

## Materials and Methods

### Cloning

Enzymes were purchased from Thermo Scientific or New England Biolabs and used according to manufacturer’s instructions. Oligonucleotides and codon-optimised synthetic genes were obtained from Integrated DNA Technologies (Coralville, Iowa, USA). All constructs were verified by sequencing (Eurofins Genomics, DE).

The mCpol-encoding genes from *Cylindrospermum stagnale* PCC 7417 (*Cst* mCpol; WP_015209627.1) and *Sphingomonas* sp. (WP_230771556.1) were cloned into pEHisV5TEV [25] for expression with a cleavable N-terminal His_8_-tag. Active site residue variants were obtained by primer-directed mutagenesis [44]. Cst CARF-TM (WP_015209626.1) and Cst mCpol were cloned into a modified pRSFDuet-based plasmid by restriction assembly to yield an N-terminal His6-TEV-CARF-TM and an untagged Cst mCpol. The expression construct for soluble *Cst* CARF was obtained by PCR-amplification of the CARF domain, followed by restriction digest and ligation into 5’-NdeI and 3’-XhoI sites of pEV5hisTEV [25]. This allows expression of *Cst* CARF with a native N-terminus and the LEHHHHHH affinity tag following residue T207 of *Cst* CARF. The *Cst* mCpol gene was cloned into the 5’-NdeI, 3’-XhoI sites of pRAT [45] to allow co-expression of the native *Cst* mCpol from an arabinose-inducible promoter with IPTG-inducible *Cst* CARF-CHis.

For plasmid challenge and phage assays, *Cst* mCpol was cloned into the 5’-NdeI, 3’-XhoI sites of pACE (Geneva Biotech, Genève, CH) and *Cst* CARF-TM was inserted into MCS1 of pRATDuet [13] to allow induction with IPTG and L-arabinose, respectively. *Cst* CARF was obtained by primer-directed mutagenesis to introduce a stop codon after residue T207 of the native sequence. The ring nuclease Crn4 from *Actinomyces procaprae* [28] and the cyclic nucleotide sponge Acb2 from *Pseudomonas* phage PaMx33 [14] were each cloned into MCS2 of pRATDuet alongside *Cst* CARF-TM in MCS1. pRATDuet-NucC has been previously described [46].

Untagged *Cst* CARF-TM and CARF constructs used in microscopy and *in vivo* fluorescence studies were obtained through restriction digest and ligation. Full length *Cst* CARF-TM was ligated into the 5’-NdeI, 3’-XhoI sites of pEHisV5TEV, which removed the N-terminal affinity tag, to give pE-*Cst* CARF-TM. The truncated *Cst* CARF was obtained by primer-directed mutagenesis to introduce a stop codon after residue T207 of the native sequence to give pE-*Cst* CARF.

### Protein Purification

#### mCpol

Cst and Sph mCpol were expressed in C43 (DE3) E. coli. 2 litres of culture were induced with 0.4 mM isopropyl-b-D-1-thiogalactoside (IPTG) at an OD_600_ of ∼0.8 and grown overnight at 16 °C. Cells were harvested (4000 rpm, 10 min, 4 °C, Avanti JXN-26, Beckman Coulter, JLA-8.1 rotor) and resuspended in lysis buffer (50 mM Tris-HCl pH 7.5, 0.5 M NaCl, 10 mM imidazole,10% glycerol, lysozyme, protease inhibitors (cOmplete™ EDTA-free protease inhibitor, Roche) and lysed by sonicating six times 1 min on ice with 1 min rest intervals (Soniprep 150, MSE). Lysates were clarified by ultracentrifugation (40,000 rpm, 30 min, 4 °C Optima L-90K, 70Ti rotor) and filtered. Proteins were purified with an immobilised metal affinity chromatography (IMAC) column (HisTrapFF crude, Cytiva), washed with 20 column volumes (CV) of buffer containing 50 mM Tris-HCl pH 7.5, 0.5 M NaCl, 30 mM imidazole and 10% glycerol, followed by a step elution with buffer containing 50 mM Tris-HCl pH 7.5, 0.5 M NaCl, 0.5 M imidazole and 10% glycerol on an NGC Chromatography System (Biorad). Protein containing fractions were concentrated and the affinity tag was removed by incubation of protein with Tobacco Etch Virus (TEV) protease (10:1) overnight at room temperature. Cleaved proteins were isolated from TEV by repeating the IMAC step and the unbound fraction collected. Size exclusion chromatography (Superdex 200 16/60, Cytiva) was used to further purify the proteins, with proteins eluted isocratically with buffer containing 20 mM Tris-HCl pH 7.5, 250 mM NaCl. The proteins were concentrated using a centrifugal concentrator (Amicon Ultra-15, Merck), aliquoted and stored frozen at -70°C.

#### Cst CARF-TM and CARF soluble domain

The Cst CARF-TM protein was transformed into competent E. coli BL21(DE3)pLysS cells. The protein was expressed by growing cells to an OD_600_ of 0.6-0.8 in LB broth at 37 ºC and inducing with 0.5 mM IPTG subsequent to a 4ºC cold shock for . Expression was then allowed to continue overnight at 16 ºC. Cells were harvested by centrifugation at 6240 x g for 15 min. Cells were resuspended in a buffer of 50 mM HEPES pH 7.3, 300 mM NaCl and supplemented with cOmplete™ EDTA-free protease inhibitor (Roche). Cells were lysed by 3 passes through a continuous flow cell disruptor (Constant Systems) at 30 kpsi. Insoluble cellular material was pelleted by centrifugation at 20,000 x g for 15 min, the resulting supernatant was centrifuged at 110,000 x g for 45 min to pellet membrane material. Membranes were homogenized in a solution of 50 mM HEPES pH 7.3, 300 mM NaCl, 20 mM Imidazole, 20 mM DDM and 0.5 mM Tris-(2-Carboxyethyl)phosphine (TCEP), then allowed to solubilise for 4 h at 4 ºC. Insoluble material was subsequently removed by centrifugation at 110,000 x g for 45 min. Cst CARF-TM was purified by TALON affinity chromatography using a wash of 20-70 mM imidazole followed by elution at 500 mM imidazole. Size exclusion chromatography on an S200 column in a buffer of 50 mM HEPES pH 7.3, 150 mM NaCl, 0.5mM DDM, 0.5 mM TCEP was carried out as an additional purification step.

The *Cst* CARF construct was co-transformed in *E. coli* C43(DE3) alongside untagged *Cst* mCpol in pRAT [45] and a single colony used to inoculate LB medium containing 50 µg/ml kanamycin and 12 µg/ml tetracycline for overnight growth at 37 °C. The overnight culture was diluted 20-fold into fresh LB medium containing antibiotics and incubated at 37 °C with shaking until the OD_600_ reached 0.6 – 0.8. *Cst* CARF production was induced with 0.1 mM IPTG and mCpol with 0.1% L-arabinose. Incubation was continued at 28 °C for 4 h, when cells were collected and the pellet stored at -20 °C. Cells were resuspended in lysis buffer (50 mM Tris, 1 M NaCl, 20 mM imidazole, 10 % glycerol, pH 7.7), lysed by sonication and the lysate was cleared by centrifugation (Ti70 rotor, 40,000 rpm, 45 min, 4 °C). *Cst* CARF was pulled down by IMAC (HisTrap™ crude, Cytiva), washed with 4% elution buffer (lysis buffer containing 0.5 M imidazole) and eluted in a gradient to 100% elution buffer. Protein-containing fractions were pooled, concentrated using an Amicon Ultra-15 (Merck) spin filter with 10 kDa MWCO, and further purified by SEC (HiLoad™ 16/600 Superdex™ 200 gp, Cytiva) in 20 mM HEPES, 250 mM NaCl, 10 % glycerol, pH 7.5. Protein-containing fractions were pooled, concentrated as before, flash-frozen and stored at -70 °C. Protein concentrations were determined by UV quantitation (NanoDrop 2000, Thermo Scientific) using calculated extinction coefficients (Benchling Biology Software, 2024).

#### Crn4 ring nuclease

Production and purification of Crn4 (Crn4a, WP_136192672, from *Actinomyces procaprae*) has been described previously [28].

### mCpol Activity

Nucleotide cyclase activity was assayed in 12.5 mM Tris-HCl, pH 8.0, 125 mM NaCl, 10 mM MgCl_2_ containing 500 µM ATP or ATP analogue and 5– 25µM (dimer) *Cst* mCpol wild type or mutant. The reaction was incubated for 3 h at 37 °C, unless stated otherwise. Enzymes were removed by ultracentrifugation (3 kDa MWCO) or by heat deactivation (95 °C for 5 min) followed by centrifugation (16,000 x g, 10 min, 4 °C). Reaction products were analysed by HPLC (Dionex UltiMate 3000 equipped with single wavelength UV detector, Thermo Scientific) as follows. Compounds from *Cst* mCpol reactions were separated on a Synergi Fusion RP column (2.5 µm, 2 x 50 mm, Phenomenex) at 0.3 ml min^-1^ and 40 °C column temperature with a linear gradient of 2 – 50 % methanol against 20 mM ammonium bicarbonate over 4.8 min and 0.2 min gradient delay. Elution was monitored by UV at 260 nm. Data were analysed using Chromeleon and visualised in Prism 10 (GraphPad). Cyclic nucleotides were identified by comparison to and co-injection with synthetic standards (Biolog, DE) or by mass spectrometry.

Selected samples were analysed by liquid chromatography-mass spectrometry on an LCQ Fleet mass spectrometer coupled to an Ultimate 3000 UHPLC module (Thermo Scientific) using the same liquid chromatography conditions as above. Electrospray ionisation was monitored in positive ion mode without fragmentation. Data were analysed with Excalibur Qual Browser software and visualised in Prism 10.

### cA_**3**_ production rate

cA_3_ production by *Sph* and *Cst* mCpol enzymes was determined using the EnzChek™ Pyrosphosphate Assay kit (Invitrogen, Fisher Scientific). Reactions contained 12.5 mM Tris-HCl, pH 8.0, 100 mM NaCl, 10 mM MgCl_2_, 0.2 mM 2-amino-6-mercapto-7-methylpurine ribonucleoside (MESG), 1 U ml^-1^ purine nucleoside phosphorylase, 0.1 U ml^-1^ pyrophosphatase, 0.5 mM ATP (Fermentas, Fisher Scientific) and 2.5 – 5 µM mCpol dimer. Background hydrolysis was determined by omitting mCpol. Pyrophosphate (1 – 10 µM) in the reaction mixture without ATP and mCpol were used to obtain the standard curve. The reaction was started by the addition of ATP or pyrophosphate standards and the absorption at 360 nm was monitored at 37 °C using a FluoStar Omega (BMG Labtech) plate reader. Data analysis was performed in Prism 10 (Graphpad). Rates were determined by linear regression, corrected for enzyme concentration with the assumption that for each cA_3_ molecule, three molecules of pyrophosphate are generated, and that non-production (pyro)phosphate release is negligible. The reported rate was the mean from three independent experiments.

### Dynamic Light Scattering

DLS measurements were performed with the Zetasizer Nano S90 (Malvern) instrument. Samples contained 60 – 100 µM protein dimer in 20 mM Tris-HCl, 75 mM NaCl, 10 mM MgCl_2_ and 0.1 % DDM as required. After centrifugation at 12,000 x g for 10 min at room temperature, the sample was centrifuged and loaded into a quartz cuvette (ZMV1012). Measurements were performed at 25 °C with three measurements of thirteen runs.

### Electrophoretic Mobility Shift Assay

[α-^32^P]-Radiolabelled cA_3_ was prepared using 45 µM *Cst* mCpol dimer in 12.5 mM Tris-HCl, pH 8.0, 100 mM NaCl, 10 mM MgCl_2_ and 300 µM [^32^P]-ATP (0.15 MBq). The reaction was incubated at 37 °C for 3 h, protein was removed by phenol-chloroform extraction and excess phenol was removed by an additional chloroform extraction step. A typical EMSA assay contained 100 nM [α-^32^P]-cA_3_ and 0 – 4 µM His_6_-CARF-TM monomer in 30 mM HEPES, pH 7.5, 90 mM NaCl, 10 mM MgCl_2_, 10 % glycerol, 0.4 mM DDM or 100 nM [α-^32^P]-cA_3_ and 0 – 14 µM CARF-CHis_6_ monomer in 12.5 mM Tris-HCl, pH 8.0, 200 mM NaCl, 10 mM MgCl_2_, 10 % glycerol, unless stated otherwise. The mixture was incubated at room temperature for 30 min before addition of G-250 Native Gel Sample Loading Buffer (Invitrogen) to 0.005% final concentration. Samples were immediately loaded onto a pre-run 6% acrylamide gel (29:1 acrylamide:bis-acrylamide). Radiolabelled material was separated for 2 h at 200 V in 1X TBE buffer and visualised by exposure to a phosphor storage screen and phosphorimaging (Cytiva).

### Protein Crosslinking

The cross-linking reaction was carried out in 20 mM HEPES, pH 7.5, 250 mM NaCl, 10% glycerol with 20 µM *Cst* CARF dimer in the absence or presence of 2.5 – 40 µM Crn4. The reaction was incubated at 37 °C for 20 min. The BS-3 (bis(sulfosuccinimidyl)suberate-d_0_, Thermo Scientific) crosslinker was then added to 0.2 mM final concentration and the mixture was incubated with gentle agitation for 30 min at 20 °C. SDS-PAGE sample loading buffer was added to each reaction and the samples were heated to 95 °C for 2 min before analysis by SDS-PAGE.

### In vitro Fluorescence Assays

Sypro Orange was purchased as a 5000X stock in DMSO (Invitrogen) and used at 5X final concentration in 20 mM HEPES, 250 mM NaCl, 10% glycerol, pH 7.5 with 9 µM CARF dimer and 3 µM Crn4 dimer wild type or [H15A] variant. The reaction was preincubated with fluorescence monitoring (ex/em 485/580 nm) at 37 °C in a FluoStar Omega plate reader (BMG Labtech) for 15 min before addition of Crn4.

BODIPY™ FL L-Cystine (BFC, Invitrogen, Fisher Scientific) was dissolved in DMSO and used at 1 µM final concentration in the same buffer as for Sypro Orange with 0.5 µM CARF and 1 µM Crn4. The reaction was preincubated for 3 min at 37 °C before addition of Crn4 with fluorescence monitoring at ex/em 485/520 nm. Data were analyzed using Prism 10.

### Plasmid Challenge Assay

The plasmid challenge assay was performed as described in [46] with the following modifications. Competent cells were prepared from individual colonies of *E. coli* C43(DE3) transformed with pACE-*Cst* mCpol, pACE-*Cst* mCpol[D63N/D64N] or pACE empty vector. pRATDuet, pRATDuet-*Cst* CARF-TM, pRATDuet-*Cst* CARF, pRATDuet-*Cst* CARF-TM-Crn4 or pRATDuet-*Cst* CARF-TM-Acb2 were used for transformation. Selective conditions were LB agar containing 100 µg ml^-1^ ampicillin for the acceptor cells only, containing ampicillin and 12 µg ml^-1^ tetracycline for uninduced transformants. *Cst* mCpol and *Cst* CARF-TM ± Crn4 or Acb2 expression was induced with 100 µM IPTG and 0.0125% or 0.05% L-arabinose, respectively. Plates were incubated overnight at 37 °C.

### Microscopy

E. coli C43(DE3) was transformed with empty vector, pE-*Cst* CARF-TM or pE-*Cst* CARF. Single colonies were used to inoculate LB containing 50 µg ml^-1^ kanamycin. After culturing overnight at 37 °C with shaking, the cultures were diluted 100-fold into fresh selective medium. Cells were grown at 37 °C to an OD_600_ of 0.5 – 0.8 and induced with 0.1 mM IPTG. Incubation was continued at 37 °C. Samples (0.5 ml) were taken before induction or 1 h after induction. Cells were collected by centrifugation at room temperature (2 min at 1,500 × g) and the pellet was washed once with PBS buffer. The pellet was resuspended in PBS buffer to an approximate OD_600_ of 5 – 10. Nile Red (Merck) was added from a 2 mg ml^-1^ stock solution in DMSO to a final concentration of 20 µg ml^-1^ and incubated in a warm environment for 30 – 45 min. Syto 9 (Invitrogen) and propidium iodide (Merck) were added to 2.5 µM and 20 µg ml^-1^ final concentration, respectively, and incubated at room temperature for 15 – 30 min.

Samples (4 µl) were applied to a thin agarose slab (1% agarose in water) applied to a standard glass microscope slide and allowed to dry for a few minutes before covering with a coverslip. Images were acquired with a fluorescence confocal microscope (DeltaVision Imaging System), connected to a digital camera. The system was conFig.d for both fluorescence and transmitted light DIC imaging. Images were recorded with an Olympus 100X/1.40 oil immersion objective lens, deconvolved using softWoRx Explorer 1.3 and further processed with Fiji image analysis software [47].

### Resazurin Assays

The same strains were used as for microscopy. Single colonies were used to inoculate LB containing 50 µg ml^-1^ kanamycin. After culturing overnight at 37 °C with shaking, the cultures were diluted 100-fold into fresh selective medium. Cells were grown at 37 °C to an OD_600_ of 0.5 – 0.8 and induced with 0.1 mM IPTG. Incubation was continued at 37 °C. Samples were taken before induction or 15 min, 30 min and 1 h after induction. The cell density was adjusted to OD_600_ 0.5 with LB medium and added to resazurin in LB (10 µg ml^-1^ final concentration) or to resazurin and polymyxin B in LB (both at 10 µg ml^-1^ final concentration) in a black, non-binding, PS 96-microtitre plate (F-bottom, Greiner). The production of resorufin was followed at 37 °C with excitation and emission wavelengths of 540 and 580 nm, respectively. Regression analysis was performed in Prism 10 (Graphpad).

## Supporting Information

**Fig. S1.**
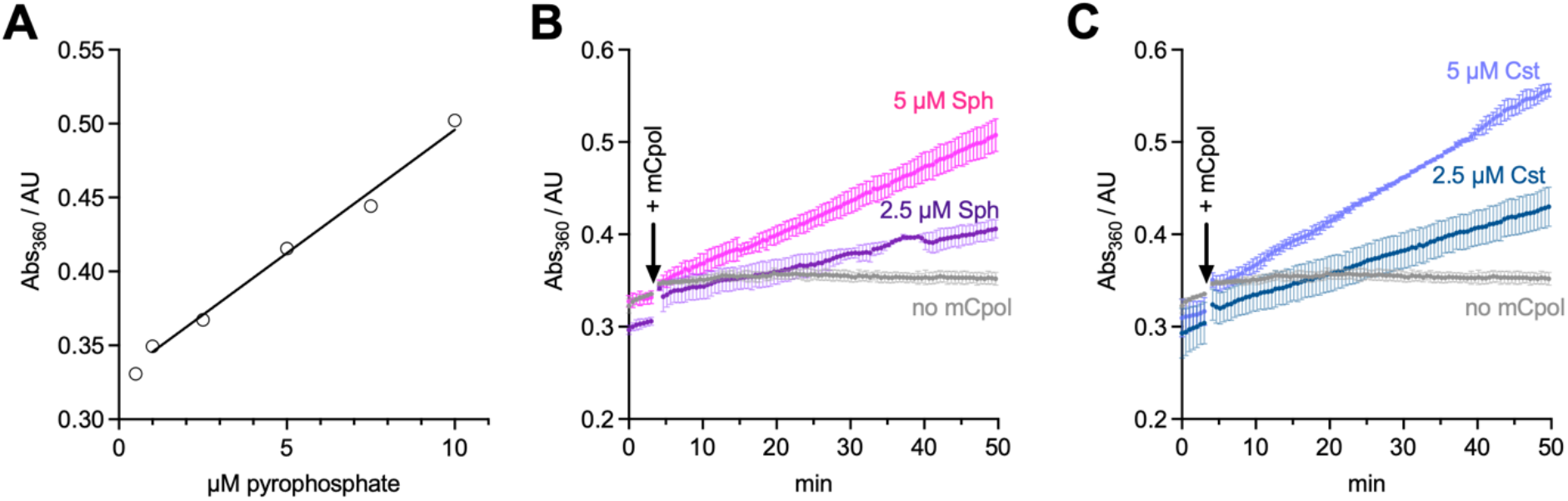
Colorimetric pyrophosphate release assays for mCpol activity. Graphs for one representative experiment are shown. **A:** Standard curve. **B:** Progress curves for Sph mCpol. **C:** Progress curves for Cst mCpol. The turnover was calculated based on the assumption that one cA_3_ molecule is produced per three pyrophosphates and that non-productive pyrophosphate release was negligible.

**Fig. S2.**
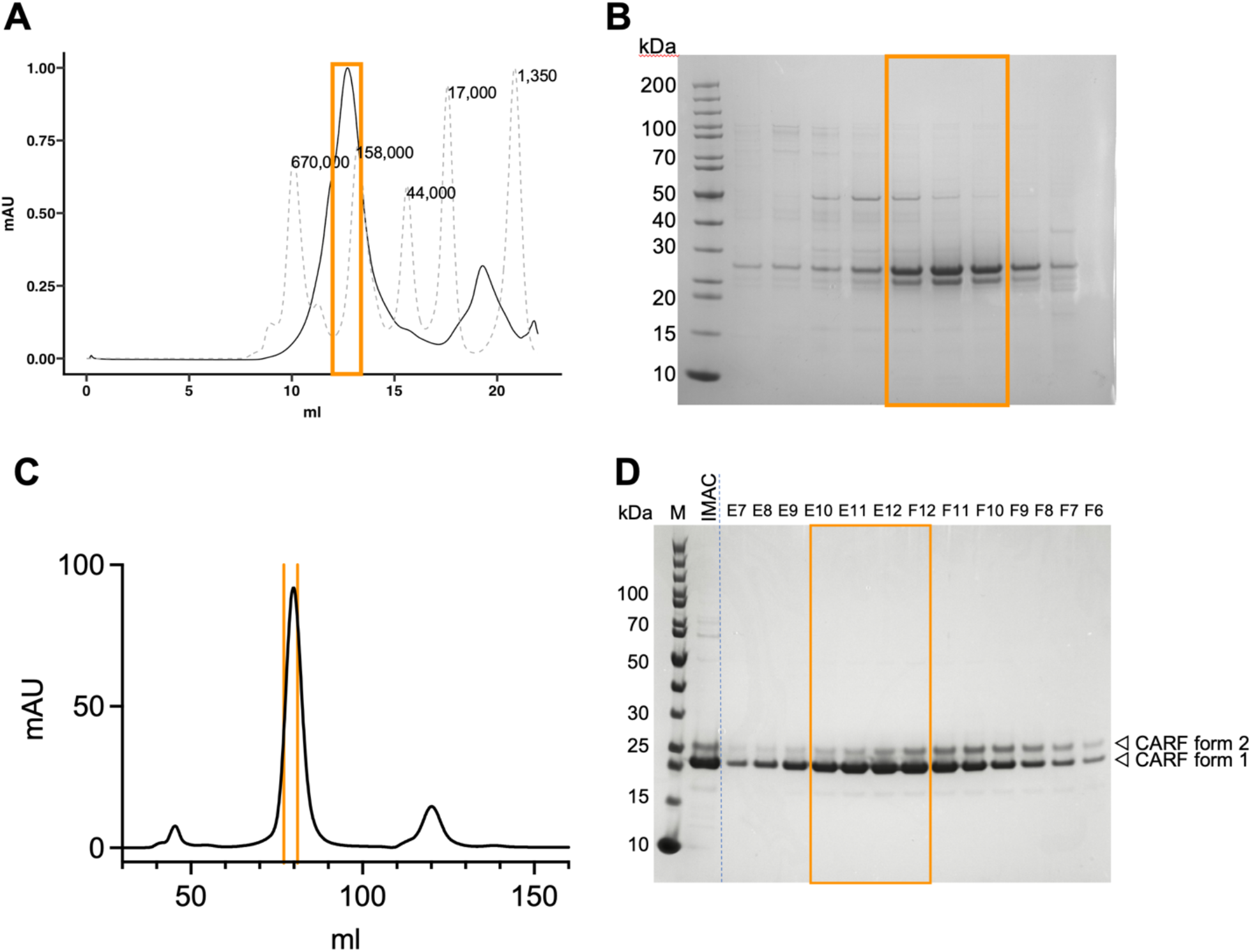
Purification of full length Cst CARF-TM (A, B) and truncated Cst CARF (C, D) effectors. Chromatograms for size exclusion chromatography (A, C) are shown and SDS-PAGE analysis of selected fractions (B, D). Pooled fractions are indicated by an orange box.

**Fig. S3.**
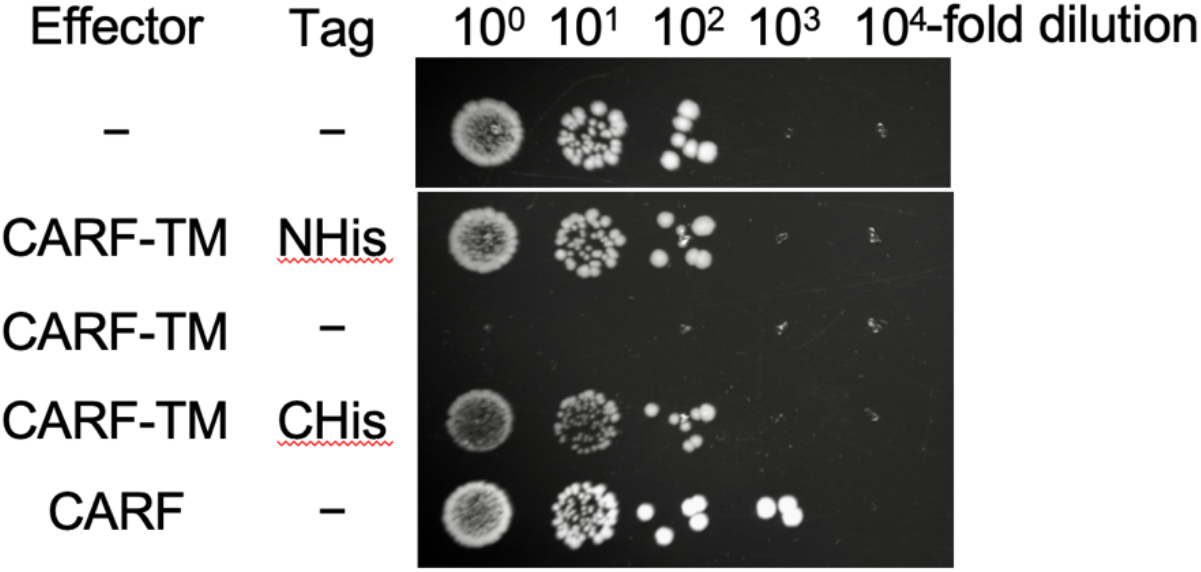
N- or C-terminal His-tags abolish CARF-TM function *in vivo*. Expression of untagged CARF-TM in E. coli prevented colony growth, but this effect was abolished when an N- or C-terminal polyhistidine tag was present.

**Fig. S4.**
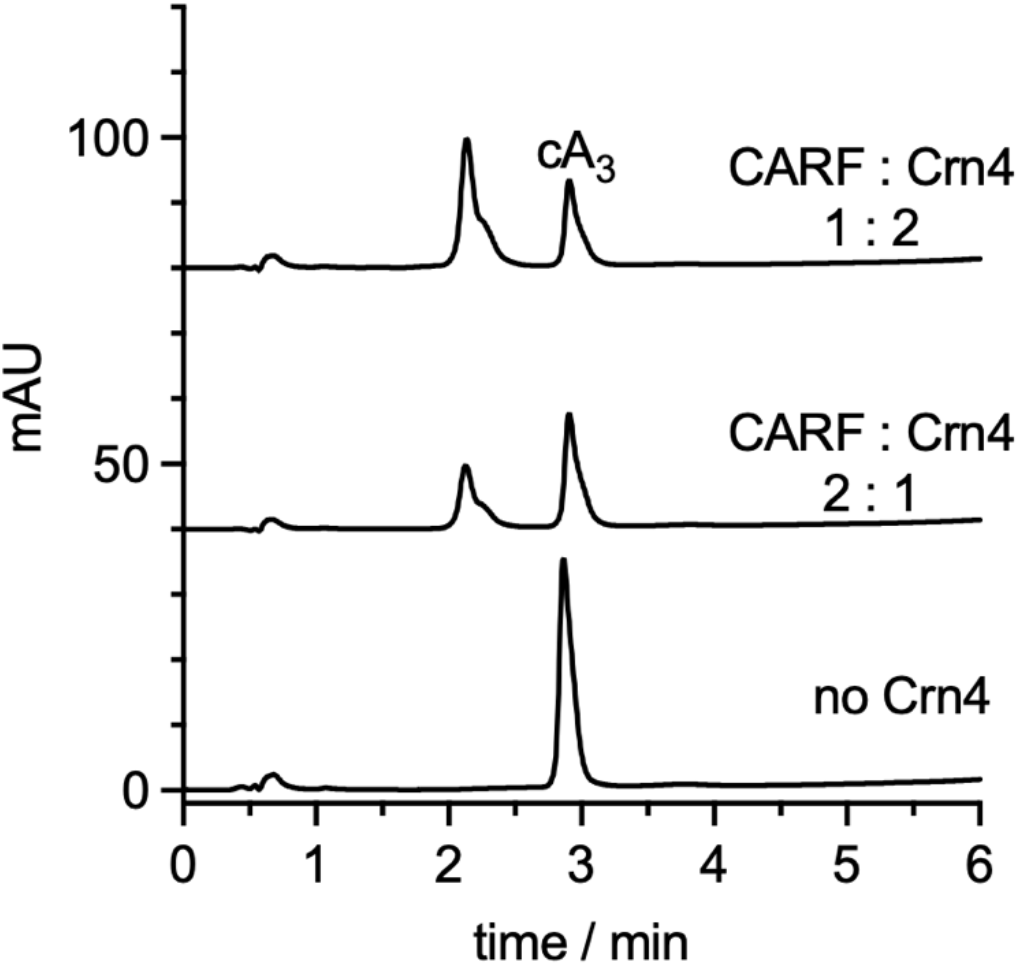
HPLC analysis of supernatant from heat denatured CARF : Crn4 mixture after 30 min incubation at 37 °C. The ring nuclease Crn4 can degrade CARF-bound cA_3_. The molar ratio of CARF to Crn4 is indicated.

**Fig. S5.**
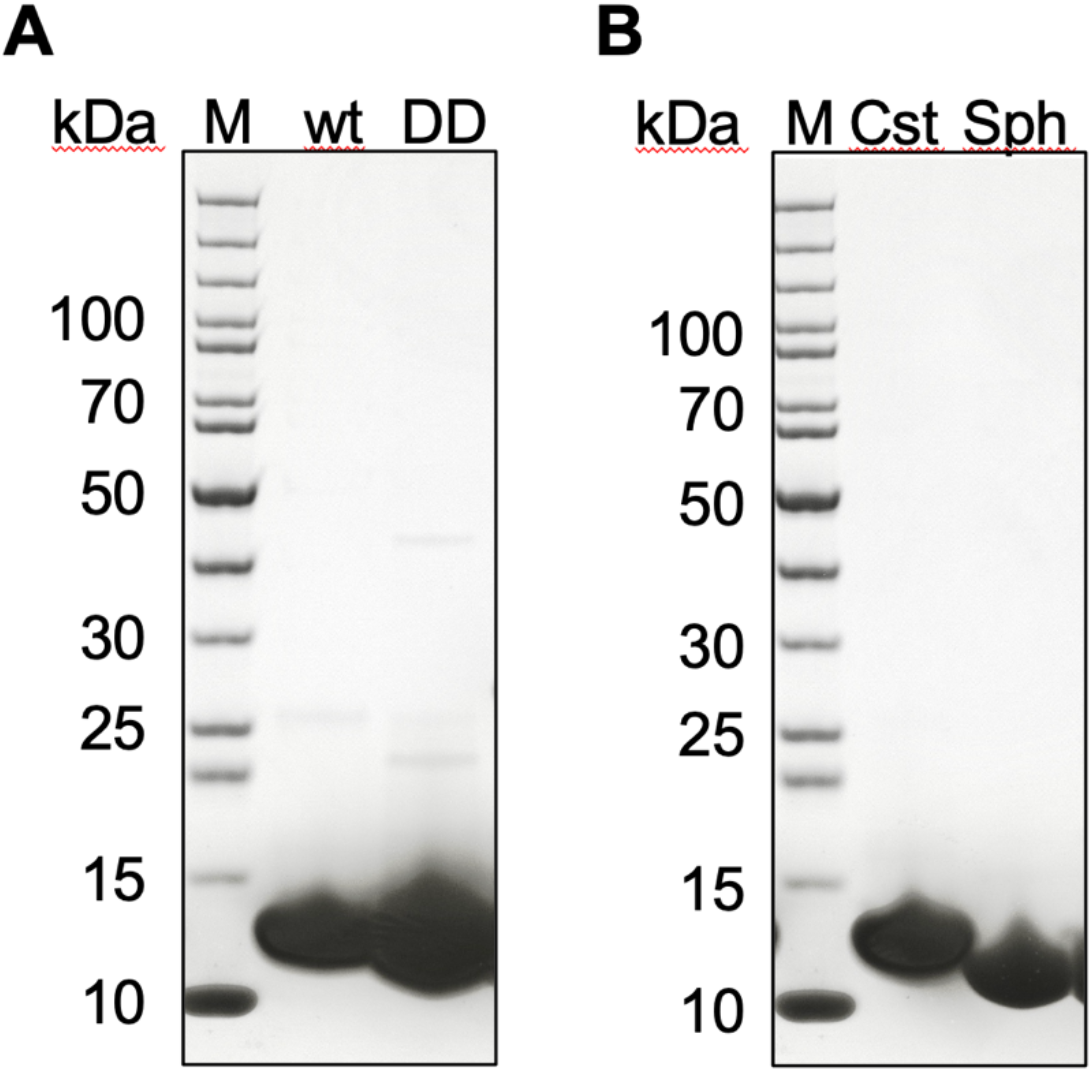
SDS-PAGE analysis of wt and variant mCpol. **A:** Cst mCpol wildtype (wt) and D63N, D64N variant (DD). **B:** Purified wild type Cst and Sph mCpol.

## References

1. Clabby T, Tesson F, Gaborieau B, Bernheim A. Why do bacteria accumulate antiphage defence systems? Philos Trans R Soc Lond B Biol Sci. 2025;380(1934):20240082. Epub 20250904. doi: 10.1098/rstb.2024.0082. PubMed PMID: 40904109.

2. Bernheim A, Sorek R. The pan-immune system of bacteria: antiviral defence as a community resource. Nature reviews. 2020;18(2):113–9. Epub 2019/11/07. doi: 10.1038/s41579-019-0278-2. PubMed PMID: 31695182.

3. Hobbs SJ, Kranzusch PJ. Nucleotide Immune Signaling in CBASS, Pycsar, Thoeris, and CRISPR Antiphage Defense. Annu Rev Microbiol. 2024;78(1):255–76. Epub 20241107. doi: 10.1146/annurev-micro-041222-024843. PubMed PMID: 39083849.

4. Slavik KM, Kranzusch PJ. CBASS to cGAS-STING: The Origins and Mechanisms of Nucleotide Second Messenger Immune Signaling. Annu Rev Virol. 2023;10(1):423–53. Epub 20230628. doi: 10.1146/annurevvirology-111821-115636. PubMed PMID: 37380187.

5. Sun L, Wu J, Du F, Chen X, Chen ZJ. Cyclic GMP-AMP synthase is a cytosolic DNA sensor that activates the type I interferon pathway. Science. 2013;339(6121):786–91. Epub 2012/12/22. doi: 10.1126/science.1232458. PubMed PMID: 23258413.

6. Duncan-Lowey B, Kranzusch PJ. CBASS phage defense and evolution of antiviral nucleotide signaling. Current opinion in immunology. 2022;74:156–63. Epub 20220202. doi: 10.1016/j.coi.2022.01.002. PubMed PMID: 35123147.

7. Tal N, Morehouse BR, Millman A, Stokar-Avihail A, Avraham C, Fedorenko T, et al. Cyclic CMP and cyclic UMP mediate bacterial immunity against phages. Cell. 2021;184(23):5728–39 e16. Epub 2021/10/14. doi: 10.1016/j.cell.2021.09.031. PubMed PMID: 34644530.

8. Kazlauskiene M, Kostiuk G, Venclovas C, Tamulaitis G, Siksnys V. A cyclic oligonucleotide signaling pathway in type III CRISPR-Cas systems. Science. 2017;357(6351):605–9. Epub 2017/07/01. doi: 10.1126/science.aao0100. PubMed PMID: 28663439.

9. Niewoehner O, Garcia-Doval C, Rostol JT, Berk C, Schwede F, Bigler L, et al. Type III CRISPR-Cas systems produce cyclic oligoadenylate second messengers. Nature. 2017;548(7669):543–8. Epub 2017/07/20. doi: 10.1038/nature23467. PubMed PMID: 28722012.

10. Rouillon C, Athukoralage JS, Graham S, Gruschow S, White MF. Control of cyclic oligoadenylate synthesis in a type III CRISPR system. eLife. 2018;7:e36734. Epub 2018/07/03. doi: 10.7554/eLife.36734. PubMed PMID: 29963983.

11. Eaglesham JB, Pan YD, Kupper TS, Kranzusch PJ. Viral and metazoan poxins are cGAMP-specific nucleases that restrict cGAS-STING signalling. Nature. 2019;566(7743):259–63. doi: 10.1038/s41586-019-0928-6. PubMed PMID: WOS:000458503900049.

12. Hobbs SJ, Nomburg J, Doudna JA, Kranzusch PJ. Animal and bacterial viruses share conserved mechanisms of immune evasion. Cell. 2024;187(20):5530–9 e8. Epub 20240827. doi: 10.1016/j.cell.2024.07.057. PubMed PMID: 39197447.

13. Athukoralage JS, McMahon SA, Zhang C, Gruschow S, Graham S, Krupovic M, et al. An anti-CRISPR viral ring nuclease subverts type III CRISPR immunity. Nature. 2020;577(7791):572–5. Epub 2020/01/17. doi: 10.1038/s41586-019-1909-5. PubMed PMID: 31942067.

14. Cao X, Xiao Y, Huiting E, Cao X, Li D, Ren J, et al. Phage anti-CBASS protein simultaneously sequesters cyclic trinucleotides and dinucleotides. Mol Cell. 2023;84:375–85. Epub 20231209. doi: 10.1016/j.molcel.2023.11.026. PubMed PMID: 38103556.

15. Leavitt A, Yirmiya E, Amitai G, Lu A, Garb J, Herbst E, et al. Viruses inhibit TIR gcADPR signalling to overcome bacterial defence. Nature. 2022;611(7935):326–31. Epub 20220929. doi: 10.1038/s41586-022-05375-9. PubMed PMID: 36174646.

16. Hadary R, Chang RB, Bechon N, Tal N, Osterman I, Yirmiya E, et al. Functional diversity of phage sponge proteins that sequester host immune signals. bioRxiv. 2025. Epub 20250824. doi: 10.1101/2025.08.24.671296. PubMed PMID: 40894804.

17. Chang RB, Toyoda HC, Hobbs SJ, Hadary R, Richmond-Buccola D, Wein T, et al. A widespread family of viral sponge proteins reveals specific inhibition of nucleotide signals in anti-phage defense. Mol Cell. 2025;85(16):3151–65 e6. doi: 10.1016/j.molcel.2025.07.016. PubMed PMID: 40845805.

18. Stella G, Marraffini L. Type III CRISPR-Cas: beyond the Cas10 effector complex. Trends Biochem Sci. 2024;49(1):28–37. Epub 20231108. doi: 10.1016/j.tibs.2023.10.006. PubMed PMID: 37949766.

19. Hoikkala V, Graham S, White MF. Bioinformatic analysis of type III CRISPR systems reveals key properties and new effector families. Nucleic Acids Res. 2024;52(12):7129–41. doi: 10.1093/nar/gkae462. PubMed PMID: 38808661.

20. Chi H, Hoikkala V, Gruschow S, Graham S, Shirran S, White MF. Antiviral type III CRISPR signalling via conjugation of ATP and SAM. Nature. 2023;622(7984):826–33. Epub 20231018. doi: 10.1038/s41586-023-06620-5. PubMed PMID: 37853119.

21. Burroughs AM, Zhang D, Schaffer DE, Iyer LM, Aravind L. Comparative genomic analyses reveal a vast, novel network of nucleotide-centric systems in biological conflicts, immunity and signaling. Nucleic Acids Res. 2015;43(22):10633–54. Epub 2015/11/22. doi: 10.1093/nar/gkv1267. PubMed PMID: 26590262.

22. Makarova KS, Timinskas A, Wolf YI, Gussow AB, Siksnys V, Venclovas C, et al. Evolutionary and functional classification of the CARF domain superfamily, key sensors in prokaryotic antivirus defense. Nucleic Acids Res. 2020;48(16):8828–47. Epub 2020/08/01. doi: 10.1093/nar/gkaa635. PubMed PMID: 32735657.

23. Sullivan AE, Nabhani A, Izrailevsky DS, Schinkel K, Hoffman CRK, Robbins LK, et al. The Panoptes system uses decoy cyclic nucleotides to defend against phage. Nature. 2025;647:988–96. Epub 20251001. doi: 10.1038/s41586-025-09557-z. PubMed PMID: 41034579.

24. Doherty EE. A minature CRISPR-Cas10 enzyme confers immunity by inhibitory signalling. Nature. 2025.

25. Rouillon C, Athukoralage JS, Graham S, Grüschow S, White MF. Investigation of the cyclic oligoadenylate signalling pathway of type III CRISPR systems. Methods Enzymol. 2019;616:191–218.

26. Grüschow S, Adamson CS, White MF. Specificity and sensitivity of an RNA targeting type III CRISPR complex coupled with a NucC endonuclease effector. Nucleic Acids Res. 2021;49:13122–34. doi: 10.1093/nar/gkab1190.

27. Lau RK, Ye Q, Birkholz EA, Berg KR, Patel L, Mathews IT, et al. Structure and Mechanism of a Cyclic Trinucleotide-Activated Bacterial Endonuclease Mediating Bacteriophage Immunity. Mol Cell. 2020;77:723–33. Epub 2020/01/15. doi: 10.1016/j.molcel.2019.12.010. PubMed PMID: 31932164.

28. Chi H, Hoikkala V, McMahon S, Graham S, Gloster T, White MF. Structure and mechanism of the broad spectrum CRISPR-associated ring nuclease Crn4. Nat Commun. 2025. doi: 10.1038/s41467-025-67607-6.

29. Abramson J, Adler J, Dunger J, Evans R, Green T, Pritzel A, et al. Accurate structure prediction of biomolecular interactions with AlphaFold 3. Nature. 2024;630(8016):493–500. Epub 20240508. doi: 10.1038/s41586-024-07487-w. PubMed PMID: 38718835.

30. Stiefel P, Schmidt-Emrich S, Maniura-Weber K, Ren Q. Critical aspects of using bacterial cell viability assays with the fluorophores SYTO9 and propidium iodide. BMC Microbiol. 2015;15:36. Epub 20150218. doi: 10.1186/s12866-015-0376-x. PubMed PMID: 25881030.

31. Pazos-Rojas LA, Cuellar-Sanchez A, Romero-Ceron AL, Rivera-Urbalejo A, Van Dillewijn P, Luna-Vital DA, et al. The Viable but Non-Culturable (VBNC) State, a Poorly Explored Aspect of Beneficial Bacteria. Microorganisms. 2023;12(1). Epub 20231225. doi: 10.3390/microorganisms12010039. PubMed PMID: 38257865.

32. Allen JL, Kennedy SJ, Shaw LN. Colorimetric assays for the rapid and high-throughput screening of antimicrobial peptide activity against diverse bacterial pathogens. Methods Enzymol. 2022;663:131–56. Epub 20211209. doi: 10.1016/bs.mie.2021.10.008. PubMed PMID: 35168786.

33. Borrelli C, Douglas EJA, Riley SMA, Lemonidi AE, Larrouy-Maumus G, Lu WJ, et al. Polymyxin B lethality requires energy-dependent outer membrane disruption. Nat Microbiol. 2025;10(11):2919–33. Epub 20250929. doi: 10.1038/s41564-025-02133-1. PubMed PMID: 41023239.

34. Nang SC, Azad MAK, Velkov T, Zhou QT, Li J. Rescuing the Last-Line Polymyxins: Achievements and Challenges. Pharmacol Rev. 2021;73(2):679–728. doi: 10.1124/pharmrev.120.000020. PubMed PMID: 33627412.

35. Limwongyut J, Moreland AS, Nie C, Read de Alaniz J, Bazan GC. Amide Moieties Modulate the Antimicrobial Activities of Conjugated Oligoelectrolytes against Gram-negative Bacteria. ChemistryOpen. 2022;11(2):e202100260. doi: 10.1002/open.202100260. PubMed PMID: 35133087.

36. Whiteley AT, Eaglesham JB, de Oliveira Mann CC, Morehouse BR, Lowey B, Nieminen EA, et al. Bacterial cGAS-like enzymes synthesize diverse nucleotide signals. Nature. 2019;567(7747):194–9. Epub 2019/02/23. doi: 10.1038/s41586-019-0953-5. PubMed PMID: 30787435.

37. Chang JJ, You BJ, Tien N, Wang YC, Yang CS, Hou MH, et al. Specific recognition of cyclic oligonucleotides by Cap4 for phage infection. Int J Biol Macromol. 2023;237:123656. Epub 20230214. doi: 10.1016/j.ijbiomac.2023.123656. PubMed PMID: 36796558.

38. Lowey B, Whiteley AT, Keszei AFA, Morehouse BR, Mathews IT, Antine SP, et al. CBASS Immunity Uses CARF-Related Effectors to Sense 3’-5’- and 2’-5’-Linked Cyclic Oligonucleotide Signals and Protect Bacteria from Phage Infection. Cell. 2020;182(1):38–49 e17. Epub 2020/06/17. doi: 10.1016/j.cell.2020.05.019. PubMed PMID: 32544385.

39. Mikheyeva IV, Sun J, Huang KC, Silhavy TJ. Mechanism of outer membrane destabilization by global reduction of protein content. Nat Commun. 2023;14(1):5715. Epub 20230915. doi: 10.1038/s41467-023-40396-6. PubMed PMID: 37714857.

40. Tan WB, Chng SS. How Bacteria Establish and Maintain Outer Membrane Lipid Asymmetry. Annu Rev Microbiol. 2024;78(1):553–73. Epub 20241107. doi: 10.1146/annurev-micro-032521-014507. PubMed PMID: 39270665.

41. Nikaido H, Vaara M. Molecular basis of bacterial outer membrane permeability. Microbiol Rev. 1985;49(1):1–32. doi: 10.1128/mr.49.1.1-32.1985. PubMed PMID: 2580220.

42. Casas-Rodrigo I, Vornholt T, Castiglione K, Roberts TM, Jeschek M, Ward TR, et al. Permeabilisation of the Outer Membrane of Escherichia coli for Enhanced Transport of Complex Molecules. Microb Biotechnol. 2025;18(3):e70122. doi: 10.1111/1751-7915.70122. PubMed PMID: 40059126.

43. Tan JMJ, Melamed S, Cofsky JC, Syangtan D, Hobbs SJ, Del Marmol J, et al. A DNA-gated molecular guard controls bacterial Hailong anti-phage defence. Nature. 2025;643(8072):794–800. Epub 20250430. doi: 10.1038/s41586-025-09058-z. PubMed PMID: 40306316.

44. Liu H, Naismith JH. An efficient one-step site-directed deletion, insertion, single and multiple-site plasmid mutagenesis protocol. BMC Biotechnol. 2008;8:91. Epub 2008/12/06. doi: 1472-6750-8-91 [pii] 10.1186/1472-6750-8-91. PubMed PMID: 19055817.

45. Grüschow S, Athukoralage JS, Graham S, Hoogeboom T, White MF. Cyclic oligoadenylate signalling mediates Mycobacterium tuberculosis CRISPR defence. Nucl Acids Res. 2019;47:9259–70. doi: 10.1101/667758.

46. Grüschow S, McQuarrie S, Ackermann K, McMahon S, Bode BE, Gloster TM, et al. CRISPR antiphage defence mediated by the cyclic nucleotide-binding membrane protein Csx23. Nucl Acids Res. 2024;52:2761–75. doi: 10.1093/nar/gkae167.

47. Schindelin J, Arganda-Carreras I, Frise E, Kaynig V, Longair M, Pietzsch T, et al. Fiji: an open-source platform for biological-image analysis. Nat Methods. 2012;9(7):676–82. Epub 2012/06/30. doi: 10.1038/nmeth.2019. PubMed PMID: 22743772.

